# Broom: Application for non-redundant storage of High Throughput Sequencing data

**DOI:** 10.1101/312306

**Authors:** Levent Albayrak, Kamil Khanipov, George Golovko, Yuriy Fofanov

## Abstract

**Motivation:** The data generation capabilities of High Throughput Sequencing (HTS) instruments have exponentially increased over the last few years, while the cost of sequencing has dramatically decreased allowing this technology to become widely used in biomedical studies. For small labs and individual researchers, however, storage and transfer of large amounts of HTS data present a significant challenge. The recent trends in increased sequencing quality and genome coverage can be used to reconsider HTS data storage strategies.

**Results:** We present Broom, a stand-alone application designed to select and store only high-quality sequencing reads at extremely high compression rates. Written in C++, the application accepts single and paired-end reads in FASTQ and FASTA formats and decompresses data in FASTA format.

**Availability:** C++ code available at https://scsb.utmb.edu/labgroups/fofanov/broom.asp

## 1 Introduction

Recent progress in HTS technology dramatically improved quality and volume of the data generated by sequencing instruments. This opens opportunities to use this technology in studies requiring high coverage of target genomes including detection of rare-variants, quasispecies analysis, and meta-barcoding. Such studies routinely produced datasets containing significant numbers of repeated and/or highly similar sequences (such as 16S rRNA produced in microbiome metabarcoding studies). The presented software application (Broom) is an attempt to employ these changes in the statistical properties and improved quality of sequencing, to refine HTS data storage strategy.

The basic principles utilized in the presented data storage/compression application include: (a) filtering and not storing low-quality data, which allows exclusion of quality scores of individual nucleotides; (b) excluding read headers and storing only a single copy of repeated sequences. Broom’s data compression approach includes storing only suffix differences between consecutive reads sorted alphabetically (similar to delta encoding techniques used in various technologies such as MPEG (Le Gall, 1992) and transforming data into binary format using bit-packing approaches specifically designed to take advantage of the limited alphabet of sequencing data.

## 2 Implementation

Broom’s quality control step (only applicable to FASTQ files) utilizes a user-defined minimum quality score threshold to replace all low quality bases by the unknown nucleotide symbol (“N”). Sequences which contain a high proportion of unknown nucleotides can be also excluded using a user defined threshold.

To transform data into compressed binary format, all sequences are sorted alphabetically using the linear time complexity Most Significant Digit Radix Sort (Knuth, 1998) algorithm which has been customized to take advantage of the limited 5 (A/T/C/G/N) letter alphabet. Sorting also places all repeated sequences together and makes it possible to calculate their copy numbers in linear time. To optimize the number of bytes required to store the copy numbers of repeated sequences, Broom categorizes alphabetically sorted sequences into four partitions: singletons (single copy) and replicated sequences with copy number values that fit into 1, 2 or 4 bytes.

In each partition Broom identifies and excludes common prefixes among consecutive sequences

**Figure 1.**
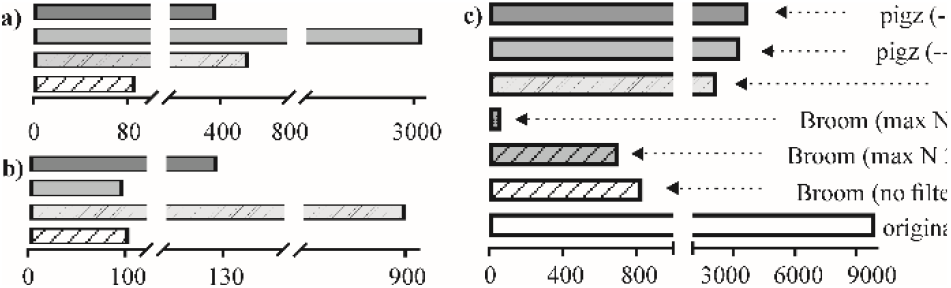
Performance comparison of Broom, pigz, and quip for SRR027520 (Illumina so that only the suffix subsequences are stored in conjunction with copy numbers, and the length of the prefix and (for paired-end and flexible length sequences) sequence lengths. Unlike the copy numbers which can be stored as contiguous blocks due to partitioning, storing other information which may vary frequently between consecutive sequences (such as prefix length, sequence length and read lengths of pairs) requires extra processing. Broom uses the most significant bit of a byte to decide whether to utilize another byte to store sequence length related information. As a result, Broom stores only one byte for length values that are up to 127 bp (7-bits) and two bytes to store length values up to 32,767 bp (15-bits). Since a de-replication step is performed prior to calculation of prefix lengths, depending on the data, the majority of the prefix values are expected to only use one byte per sequence which results in a large savings in the resulting compressed file size.

To compress suffix sequences, they are concatenated into a single string of bytes in which each nucleotide is represented as an integer value from 0 to 5. The main advantage of using a contiguous block of bytes, rather than individually allocated suffix sequences, is that it maximizes utilization of the CPU cache for further processing on the suffixes.

In the final step of the algorithm, the frequency and length span of unknown nucleotides in the suffix byte string, are analyzed to choose between two alternative compression strategies. If the unknown nucleotides appear sparsely and form islands, Broom converts unknown nucleotides into “A”s in place and performs a base 4 transformation (for the 4 letter alphabet of A, T, C and G) of the suffix bytes by converting 32 nucleotides at a time into 8-byte integer values while the location and span of the unknown nucleotides are stored separately. Alternatively, it performs a base 5 transformation of the suffix bytes. Base 5 transformation of the suffix bytes allows the storage of 27 nucleotides per 8-byte integer. Transforming sequences into base 5 in 8-byte blocks, utilizes 2.37 bits per nucleotide which is preferred over a simple 3 bit per nucleotide transformation (21 nucleotides per 8-byte block) to represent 5 possible nucleotide values. The choice of 8-byte blocks is due to the efficient processing of 64-bit integers by the CPU without the need for excessive operations required for bit packing.

Even though the design principals of Broom make it effective in compressing HTS data, it also supports compression of genome files in FASTA format, albeit less effectively than HTS datasets. The genome module performs lossless compression of FASTA files retaining sequence headers. The nucleotide sequences are compressed using base 4 or base 5 transformations while none of the sorting and prefix identification steps are performed as these steps only take advantage of properties of HTS data.

It is important to mention that if no quality filtering is applied, Broom’s memory footprint is roughly twice the size of the original file. Additionally, the presented version of Broom is strictly single-threaded and it can be run in parallel using the operating system provided utilities. All characters in the nucleotide sequences are capitalized and any nucleotide character outside of the set of A, T, C, G, and N are replaced with “N”s.

## 3 Results and Performance

The performance comparison between Broom and the latest versions of quip (Jones *et al*., 2012) (version 1.18) and pigz (Adler, 2007) (version 2.34) were made using 5 FASTQ files (ranging from 300 megabytes to 10 gigabytes), downloaded from the Sequence Read Archive (Leinonen *et al*., 2011). Three different filtration parameters were used for Broom: without discarding any reads containing unknown or low quality nucleo tides, discarding reads with unknown or low quality (minimum quality score 15) nucleotides more than 25% of the read length, and discarding reads with one or more unknown or low quality nucleotides). pigz was tested using *fast* (fastest compression method) and *best* (slowest but the best compression method) options. All calculations were performed on a 1400 MHz AMD Opteron(tm) computer with 6386 SE processor, 2048 Kb cache, and 512 GB RAM running CentOS Linux. The timing measurements were made using the Linux “time” command and CPU time was calculated by adding user and sys fields.

In all the tested cases, files generated by Broom are dramatically (up to 150 fold) smaller, especially when filtration parameters were set to exclude reads containing low quality nucleotides (Figure 1 and Supplementary documents). It is also important to emphasize that in all the tested cases Broom achieved significantly better compression/decompression times.

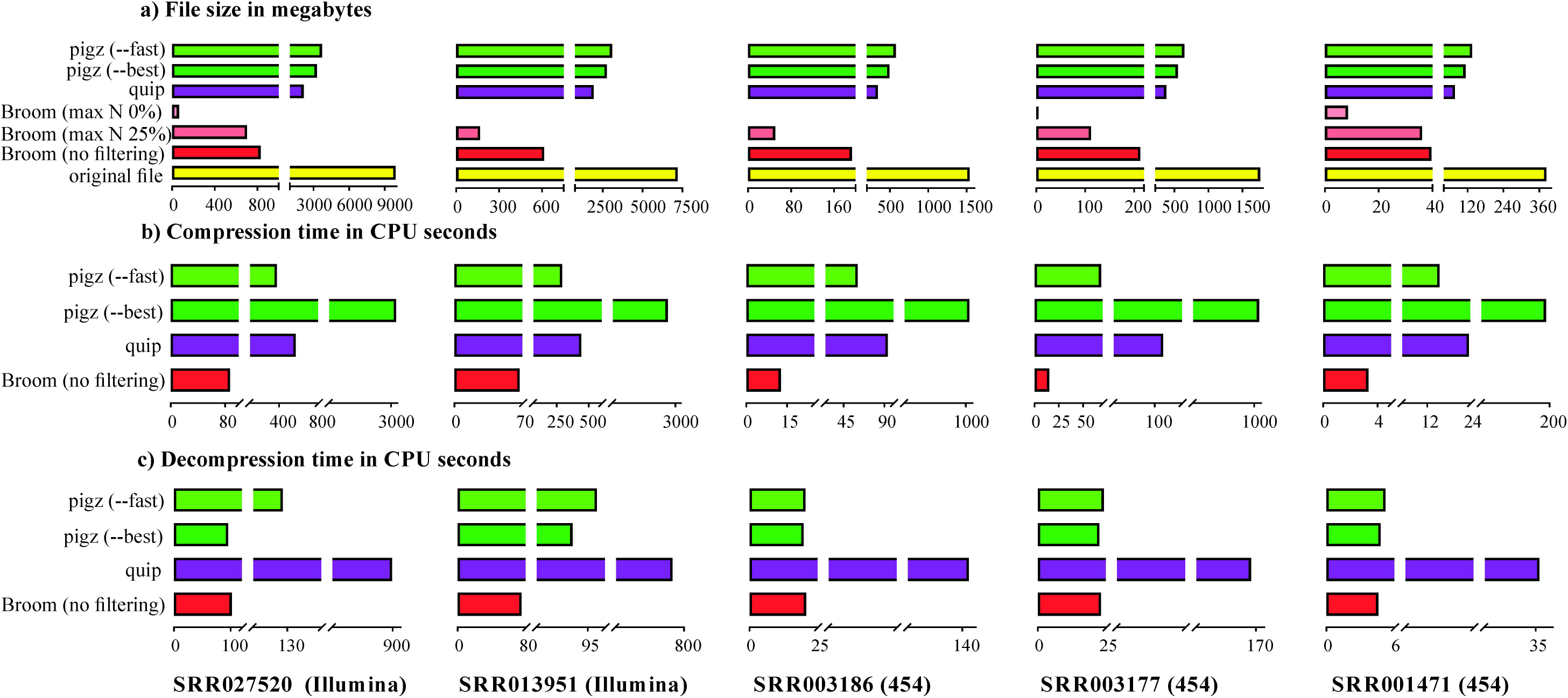

## Acknowledgements

The authors would like to thank Otto Dobretsberger for the fruitful discussions and constructive comments.

